# RNF8 is not required for histone-to-protamine exchange in spermiogenesis

**DOI:** 10.1101/2020.12.05.413005

**Authors:** Hironori Abe, Rajyalakshmi Meduri, Ziwei Li, Paul R. Andreassen, Xin Zhiguo Li, Satoshi H. Namekawa

**Affiliations:** Division of Reproductive Sciences, Cincinnati Children’s Hospital Medical Center, Cincinnati, Ohio, 45229, USA; Department of Microbiology & Molecular Genetics, University of California, Davis, California, 95616, USA; Center for RNA Biology: From Genome to Therapeutics, Department of Biochemistry and Biophysics, Department of Urology, University of Rochester Medical Center, Rochester, New York, 14642, USA; Division of Experimental Hematology & Cancer Biology, Cincinnati Children’s Hospital Medical Center, Cincinnati, Ohio, 45229, USA

**Keywords:** Sperm, Spermiogenesis, histone-to-protamine exchange, MIWI, RNF8

## Abstract

A member of the PIWI family of proteins, MIWI, binds to Piwi-interacting RNA (piRNA) and is essential in mouse spermiogenesis. A recent study demonstrated that MIWI is an essential regulator of the histone-to-protamine exchange in spermiogenesis and that this function is mediated by its binding to an ubiquitin ligase, RNF8. However, here we confirm that RNF8 is not required for histone-to-protamine exchange in spermiogenesis. We show that histone-to-protamine exchange takes place in *Rnf8*-deficient mice, while RNF8 mediates ubiquitination of H2A on the sex chromosomes in meiosis, the prior stage of spermatogenesis. Therefore, the infertile phenotype of MIWI mutant mice cannot be explained by a RNF8-mediated mechanism in spermiogenesis.

## Introduction

To prepare fertilization, male germ cells undergo a massive nuclear transformation in which nuclear histones are largely replaced with highly basic proteins, called protamines, to produce condensed sperm nuclei. The mechanism underlying histone-to-protamine exchange remains largely unknown, although several key aspects have been identified. Genome-wide histone acetylation represents an initial signal for histone-to-protamine exchange (Goudarzi et al., 2014). Following acetylation, the histone variant H2A.L.2 works with transition proteins, TNP1 and TNP2, for the loading of protamines (Barral et al., 2017). A histone-ubiquitin ligase RNF8 was reported to be required for genome-wide histone acetylation and subsequent histone-to-protamine exchange based on the absence, genome-wide, of histone acetylation, and of two protamine proteins, PRM1 and PRM2 in *Rnf8*-deficient (KO) mice (Lu et al., 2010). However, using the same line of *Rnf8*-KO mice, we showed that genome-wide accumulation of histone H4K16 acetylation and incorporation of PRM1 took place in *Rnf8*-KO elongating spermatids; the effects of RNF8 on PRM2 deposition in elongating spermatids was not examined at that time (Sin et al., 2012). In contrast, we did confirm a role for RNF8 in mediating genome-wide ubiquitination of H2A in elongating spermatids. Thus, our previous study demonstrated that RNF8-mediated ubiquitinated H2A is associated neither with H4K16 acetylation nor with the histone-to-protamine exchange in elongating spermatids.

Nonetheless, later studies have continued to postulate a requirement for RNF8 in the histone-to-protamine exchange. Among these, a recent study claimed that a mouse PIWI protein, MIWI, controls RNF8-mediated histone-to-protamine exchange through the direct regulation of RNF8 by MIWI in spermiogenesis in mice (Gou et al., 2017). PIWI proteins are evolutionarily-conserved proteins that are expressed in germ cells and are associated with Piwi-interacting RNA (piRNA) for suppression of retrotransposons (Juliano et al., 2011; Siomi et al., 2011). MIWI is one of the two PIWI proteins that is expressed during the pachytene stage in mice (Li et al., 2013) and loss of its function leads to spermiogenesis arrest (Deng and Lin, 2002). Based on the finding that MIWI regulates RNF8, Gou et al. (2017) concluded that MIWI has a piRNA-independent function, but instead has a RNF8-dependent function in histone-to-protamine exchange. Further, an additional study suggested that RNF8 is required for histone-to-protamine exchange based on the presence of testis-specific histone H2B in sperm nuclei of *Rnf8*-deficient mice (Guo et al., 2018). These studies have spawned a lingering view in the field that RNF8 is required for histone-to-protamine exchange (Wang et al., 2019; Yan, 2017).

In the present study, to clarify the function of RNF8 in histone-to-protamine exchange, we rigorously examined whether RNF8 is involved in this process in two independent laboratories at different institutions (the Namekawa and Li laboratories). Here, we demonstrate that histone-to-protamine exchange takes place in *Rnf8*-deficient mice, confirming our previous conclusion that RNF8 is not required for histone-to-protamine exchange in spermiogenesis. Instead, RNF8 has a nuclear function in the ubiquitination of the sex chromosomes during meiosis and in the activation of sex chromosome-linked genes in postmeiotic spermatids (Adams et al., 2018; Hasegawa et al., 2015; Sin et al., 2012). Therefore, we again refute the findings of Lu et al. (2010) that RNF8 is required for histone-protamine replacement. Furthermore, as a consequence, the infertility phenotype of MIWI mutant mice reported in Gou et al. (2017) cannot be explained by a RNF8-mediated mechanism in spermiogenesis, challenging the main mechanistic conclusion of that article.

## Results and Discussion

To confirm whether RNF8 is required for histone to protamine exchange, we (the Namekawa and Li laboratories) have independently examined *Rnf8*-deficient (KO) mice (Minter-Dykhouse et al., 2008). The results were consistent between our two labs, and here we present representative data. This *Rnf8*-KO mouse line was generated from an embryonic stem cell line, RRR260, in which the *Rnf8* gene was disrupted by a gene trap mutation, and the absence of RNF8 protein was previously confirmed (Santos et al., 2010). This same line of *Rnf8*-KO mice was widely used in previous studies of the function of RNF8 in spermatogenesis (Adams et al., 2018; Guo et al., 2018; Hasegawa et al., 2015; Ichijima et al., 2011; Lu et al., 2010; Lu et al., 2013; Santos et al., 2010; Sin et al., 2012; Sin et al., 2015), including the initial study by Lu et al. (2010) which suggested a requirement for RNF8 in mediating histone-to-protamine exchange.

Since Lu et al. reported that RNF8-dependent histone ubiquitination induces H4K16 acetylation as an initial step of histone removal (Lu et al., 2010), we first revisited whether H4K16 acetylation is RNF8-dependent or not. In accordance with our previous conclusion (Sin et al., 2012), we confirmed that the distribution of H4K16 acetylation was comparable between testicular sections of *Rnf8*-KO and control littermate wild-type testes (Figure 1A). This result was independently confirmed by comparable immunostaining signals of pan-acetylation on histone H4 between *Rnf8*-KO testes and wild-type controls (Figure 1B). However, RNF8 is indeed required for H2A ubiquitination on entire nuclei of elongating spermatids (Sin et al., 2012). Thus, we conclude that RNF8-dependent histone ubiquitination does not induce genome-wide H4K16 acetylation in elongating spermatids.

**Figure 1.**
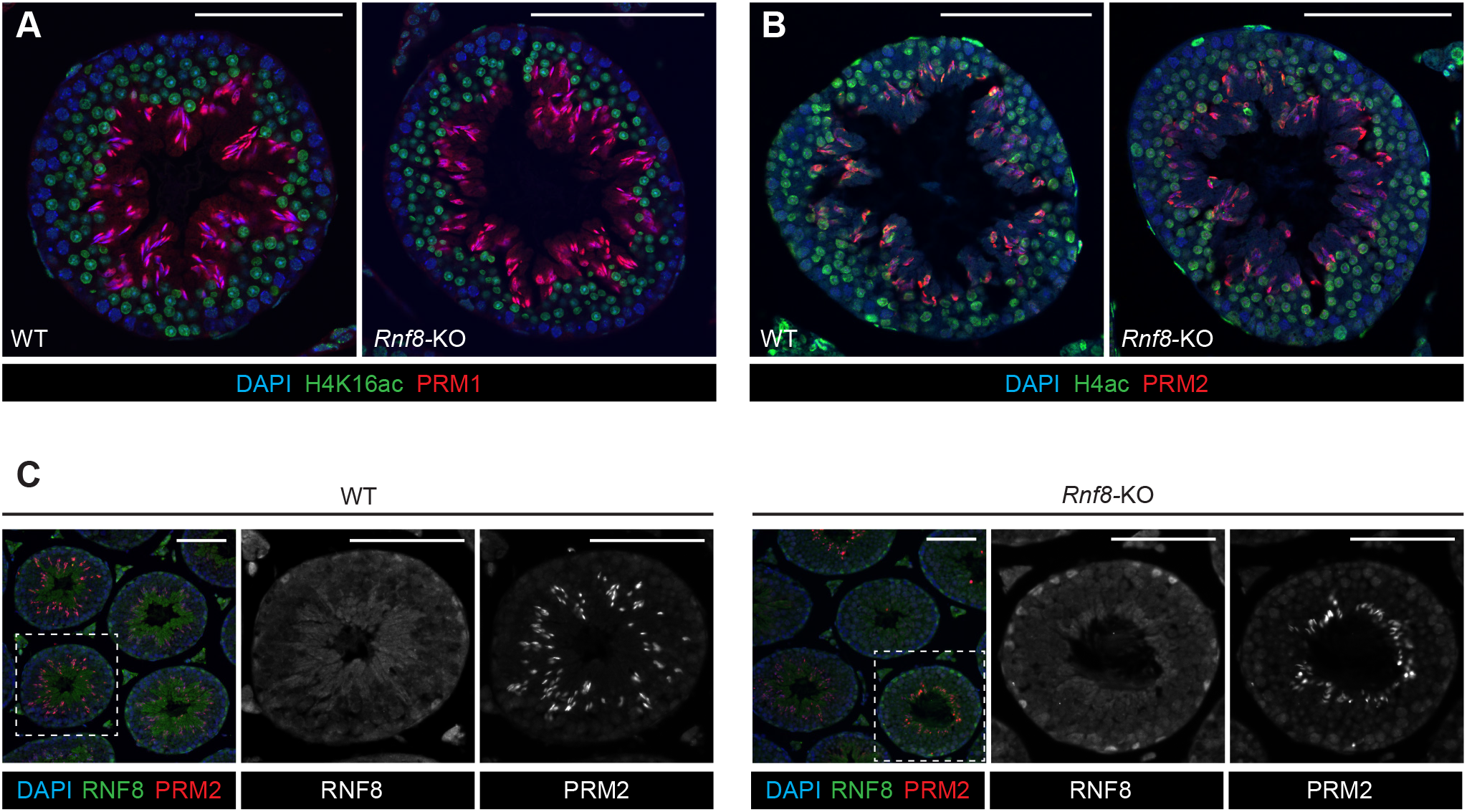
RNF8 is not required for H4K16 acetylation and histone-to-protamine exchange in spermiogenesis. (**A, B, C**) Testis sections from wild-type (WT) and *Rnf8*-KO mice immunostained with antibodies shown in the panels (H4K16ac: H4K16 acetylation; H4ac: pan-acetyl H4). Dashed squares are magnified in panels to the right. Scale bars: 100 μm.

Next, we examined the distribution of protamines (PRM1 and PRM2) on testicular sections of *Rnf8*-KO mice. We found that both PRM1 and PRM2 were present, and the distributions were comparable between *Rnf8*-KO testes and wild-type controls (Figures 1A-C). These results confirm our previous conclusion that PRM1 incorporation is independent of RNF8 (Sin et al., 2012), and in addition now shows that incorporation of two kinds of protamines (PRM1 and PRM2) takes place in spermiogenesis in *Rnf8*-KO testes.

To further confirm RNF8-independent protamine incorporation, we prepared smear slides from epididymal sperm of *Rnf8*-KO mice. *Rnf8*-KO epididymis contained sperm (Lu et al., 2010), suggesting that spermiogenic progression was not abrogated in *Rnf8*-KO testes. Further, we found that PRM1 was present in *Rnf8*-KO epididymal sperm, and the distribution of PRM1 was comparable between *Rnf8*-KO and wild-type sperm (Figure 2A). PRM2 was also present in *Rnf8*-KO epididymal sperm, and the distribution of PRM2 was comparable between *Rnf8*-KO and wild-type sperm (Figure 2B). Together, these results confirm that RNF8 is not required for the histone-to-protamine exchange.

**Figure 2.**
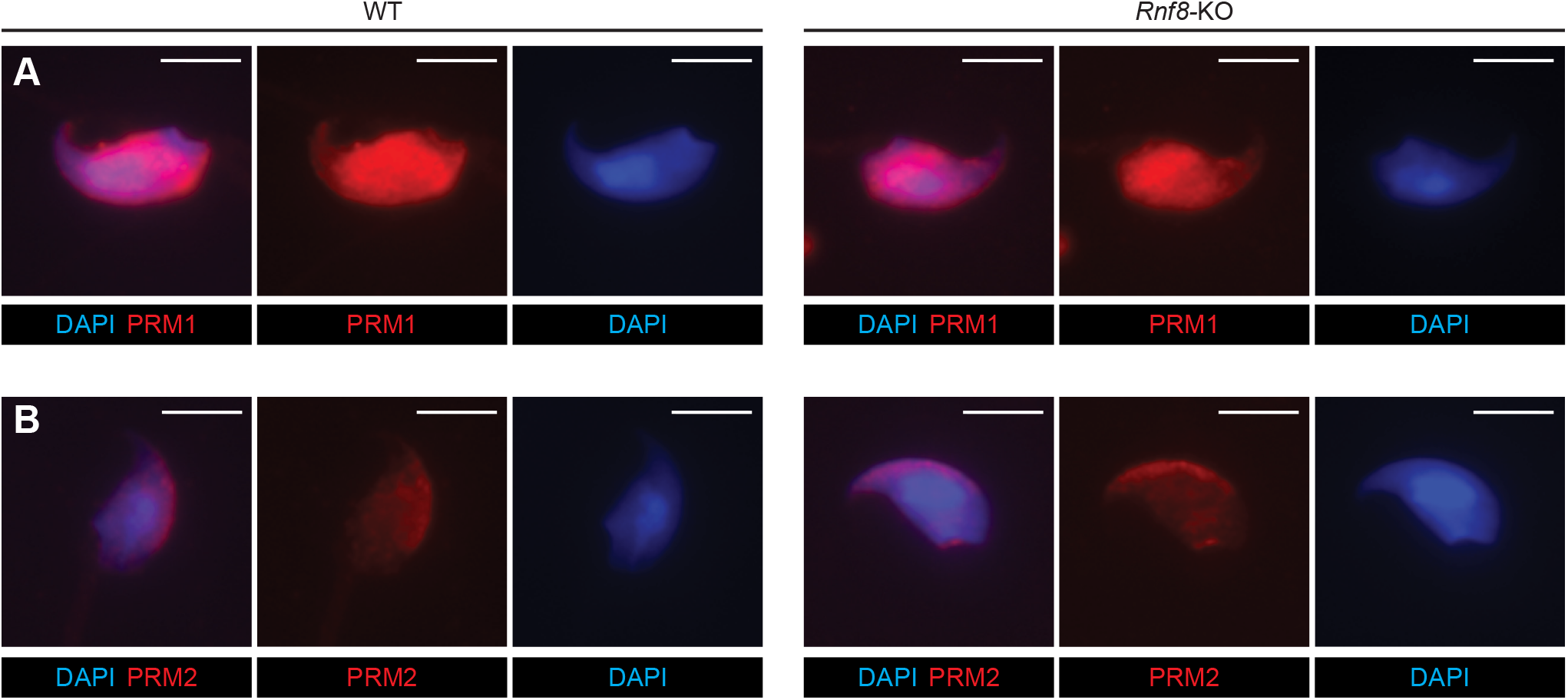
RNF8 independent incorporation of PRM1 and PRM2 in epididymal sperm. (**A, B**) Smear preparations from wild-type (WT) and *Rnf8*-KO epididymal sperm immunostained with antibodies against PRM1 or PRM2. Scale bars: 5 μm.

A possible explanation for why the initial study by Lu et al. did not observe protamine incorporation into sperm nuclei (Lu et al., 2010) is the inability of *Rnf8*-KO spermatogenesis to reach the stage of protamine incorporation in certain specific genetic backgrounds or environments. In support of this possibility, Lu et al. showed that the late stages of spermiogenesis were abrogated in the *Rnf8*-KO testis. (Lu et al., 2010). On the other hand, *Rnf8*-KO mice in our colony did not show abrogation of spermiogenic progression in *Rnf8*-KO testes, enabling us to confirm that RNF8 is not required for histone-to-protamine exchange. Curiously, the same *Rnf8*-KO mice were reported to be subfertile in another study (Guo et al., 2018). While that study claimed that RNF8 is required for histone-protamine exchange, fertile *Rnf8*-KO sperm must have underwent normal histone-protamine exchange to enable fertilization, supporting our conclusion that RNF8 is not required for the histone-to-protamine exchange. These studies together suggest that the phenotypic outcomes (spermiogenic progression) were different among the various studies due to distinct genetic backgrounds or environments, thereby leading to conflicting conclusions about the role of RNF8 in the histone-to-protamine exchange. Specifically, some factor(s) associated with a particular genetic background and/or environment may have acted as a modifier of spermiogenic progression in *Rnf8* KO mice.

In spite of the variable spermiogenic progression among studies, previous studies clearly agree that RNF8 has a function in male meiosis prior to spermiogenesis. RNF8 has a nuclear function in the regulation of sex chromosomes, and *Rnf8*-KO mice consistently show abrogation of histone ubiquitination on the sex chromosome during meiosis, as also observed by Lu al. (2010) (Adams et al., 2018; Hasegawa et al., 2015; Lu et al., 2010; Lu et al., 2013; Santos et al., 2010; Sin et al., 2012). Notably, this phenotype was observed in 100% of the meiotic cells we have examined. This complete phenotype may be due to a requirement for the DDR pathway, which includes RNF8, in the regulation of unsynapsed sex chromosomes during meiosis (Ichijima et al., 2011; Ichijima et al., 2012), and DDR-dependent coordination of meiotic progression (Abe et al., 2020). Downstream of RNF8-mediated ubiquitination on the sex chromosomes, active epigenetic modifications are established on meiotic sex chromosomes, and a portion of sex chromosome-linked genes are activated in postmeiotic spermatids (Adams et al., 2018; Hasegawa et al., 2015; Sin et al., 2012; Sin et al., 2015). Variable abnormal gene expression of sex chromosome-linked genes in *Rnf8*-KO testes could modulate phenotypic outcomes (i.e. spermiogenic progression) in distinct genetic backgrounds or environments among different studies (Guo et al., 2018; Lu et al., 2010; Sin et al., 2012). Additionally, a chromatin-remodeling nuclear protein, CHD5 has been reported as another factor that controls hyperacetylation of H4 at K5/8/16, histone-to-protamine exchange and chromatin condensation in sperm heads (Li et al., 2014; Zhuang et al., 2014), which shows substantial similarity with the *Rnf8-*KO phenotype reported by Lu et al. (2010). This is an example of a factor that could potentially be affected by differences in genetic background or environment and thereby account for variable phenotypes of the *Rnf8*-KO in particular studies.

In the studies by Gou et al. (2017), which claimed that MIWI-dependent regulation of RNF8 is critical for histone-to-protamine exchange, the authors did not directly examine *Rnf8*-KO mice. However, the authors may have miscited the literature about the function of RNF8. We already described that RNF8 is not required for histone-to-protamine exchange (Sin et al., 2012), but in Gou et al. (2017), RNF8 was described to be “responsible for histone-to-protamine exchange (Lu et al., 2010), but not critical for protamine expression (Sin et al., 2012)”. Therefore, the study by Gou et al., appeared to lack the literature-based foundation that RNF8 is responsible for histone-to-protamine exchange. Gou et al. described the phenotype of a novel MIWI mutation of the N-terminal D-box (heterozygous germline-specific knockin of the R218A/L221A mutations on a single allele of the *Miwi* gene, referred to as *Miwi*^+/*DB*^) (Gou et al., 2017). The late stage of spermiogenesis was abrogated in the testes of *Miwi*^+/*DB*^ mice (Gou et al., 2017). Although the similarity between *Miwi*^+/*DB*^ spermiogenesis and *Rnf8*-KO mice in the Lu et al. (2010) study was described (Gou et al., 2017), the most straightforward interpretation is that *Miwi*^+/*DB*^ spermiogeneis did not reach the stage of histone-to-protamine exchange, and there is no direct evidence that the phenotype of *Miwi*^+/*DB*^ spermiogenesis is due to the loss-of-function of RNF8. Instead the case is built upon the demonstration of an interaction between MIWI and RNF8, and the fact that modulation of MIWI levels has an effect on RNF8 localization to nuclei.

Further, Gou et al. examined RNF8 using a commercial anti-RNF8 antibody. However, the antibody does not specifically recognize RNF8; We detected non-specific signals in Western blots of testicular lysates and immunostaining of testicular sections using *Rnf8*-KO testes (Figure 1C and 3). In fact, there was no evidence of a specific RNF8 signal on Western blots (Figure 3). This is a key point because as noted above, the evidence for a MIWI-dependent role of RNF8 in histone-to-protamine exchange was indirect and rested in significant part on a purported role for MIWI in controlling nuclear localization of RNF8; potential utilization of a non-specific antibody for that experiment clearly calls the result and conclusion into question. Taken together, the infertile phenotype of *Miwi*^+/*DB*^ mice reported in Gou et al. (2017) cannot be explained by the RNF8-mediated mechanism in spermiogenesis, challenging the main mechanistic conclusion made by Gou et al. (2017).

**Figure 3.**
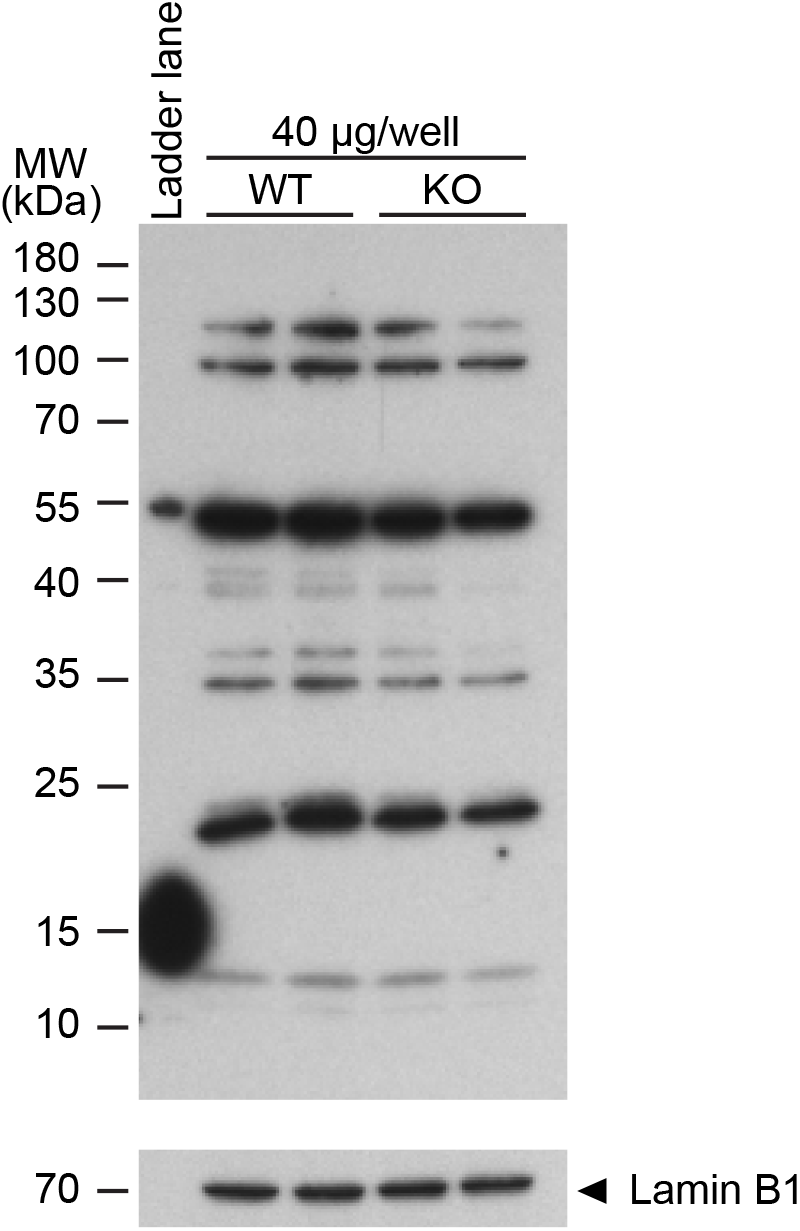
Failure of an anti-RNF8 antibody to specifically recognize RNF8. Western blot of wild-type (WT) and *Rnf8*-KO lysates from whole mouse testes probed with an anti-RNF8 antibody. 40 μg protein of sample was loaded in each lane. Two independent samples for WT and *Rnf8*-KO testes are shown. Loading control: Lamin B1. The expected molecular weight (MW) of RNF8 is 56-60 kDa. Of critical importance, the band most proximal to the expected size of RNF8 was equally present in extracts in WT and *Rnf8*-KO lysates, and no other prominent band displayed a clear difference between these genotypes.

## METHODS

### Mouse lines

The *Rnf8*-KO we utilized was generated previously from the embryonic stem cell line RRR260 (Bay Genomics) (Minter-Dykhouse et al., 2008) and is on C57BL/6 backgrounds. All subsequent experimental work was performed under protocol no. IACUC2018-0040 approved by the Institutional Animal Care and Use Committee of Cincinnati Children’s Hospital and Medical Center.

### Immunohistochemistry

For the preparation of testis paraffin blocks, excised testes were fixed with 4% paraformaldehyde in phosphate-buffered saline (PBS) containing 0.1% Triton X-100 at 4°C overnight. Testes were dehydrated and embedded in paraffin. For histological analyses, 6 μm-thick paraffin sections were deparaffinized. The sections were autoclaved in Target Retrieval Solution, Citrate pH 6.1 (DAKO, S-1700) or Antigen Unmasking Solution, Citric Acid Based (Vector, H-3300), 121°C, 100 kPa (15 psi) for 10 min. The sections were blocked with Blocking One Histo (Nacalai USA, 06349-64) at room temperature (RT) for 10 min; then, the sections were incubated with primary antibodies diluted in PBS at 4°C overnight. The following antibodies were used at the indicated dilutions [format: host anti-protein (source or company with product/catalog number), dilution]: rabbit anti-H4K14ac (Millipore, 06-762), 1/200; rabbit anti-H4ac (Millipore, 06-866), 1/200; mouse anti-Protamine 1 (Briar Patch Biosciences, Hup1N), 1/200; goat anti-Protamine 2 (Santa Cruz, sc-23104), 1/200; rabbit anti-RNF8 antibody (Proteintech, 14112-1-AP). The resulting signals were detected with appropriate secondary antibodies conjugated to Alexa 488 or 555 (ThermoFisher Scientific), diluted 1/1000 in PBS and incubated at RT for 1 h. Sections were counterstained with the DNA-binding chemical 40, 6-diamidino-2-phenylindole (DAPI; Sigma, D9542-5MG) diluted to 1 mg/mL in water for 5 min, then mounted with ProLong™ Gold Antifade Mountant (Thermo Fisher Scientific, P36930). Images were obtained with an A1RSi Inverted Confocal Microscope (Nikon) and processed with NIS-Elements Basic Research (Nikon), Photoshop (Adobe), and Illustrator (Adobe).

### Smear preparations

Sperm were obtained from cauda epididymides of WT and *Rnf8*-KO at 11 weeks of age. To make sperm suspensions, first, a drop of 400 μL of M2 medium (Sigma, M7167) was prepared on a 35 mm plastic dish and it was fully covered with heavy paraffin oil (Sigma, PX0046-1); then the dish was kept on a hot plate at 37 °C until use. The obtained cauda were placed in the paraffin oil and a drop of sperm was squeezed out from the cauda using a forcep and a needle. The sperm drop was transferred into the drop of M2 medium. After incubation at 37 °C, 5% CO2 for 30 min, sperm swimming up in M2 medium were collected using pipettes. After washing two times with PBS followed by centrifugation at 3,000 *g* for 3 min at RT, sperm pellets were resuspended into 200 μL of PBS and sperm numbers were counted. An aliquot containing 1 × 10^5^ sperm was used to make smears and slides were stored at −80°C until staining was performed.

### Immunostaining on sperm

We followed a previously described protocol (Balhorn et al., 2018). Mouse anti-Prm1 antibody (1:500 dilution; Briar Patch Biosciences, Livermore, CA, USA; Hup1N) and mouse anti-Prm2 antibody (1:200 dilution; Briar Patch Biosciences, Livermore, CA, USA; Hup2B) were used. Secondary antibodies conjugated with either Alexa Fluor 488 or 594 (Molecular Probes, Eugene, OR, USA) were used at a dilution of 1:500.

### Western blotting

Western blot experiments were replicated twice; for each experiment, all samples were run on the same gel. Detunicated testis pieces obtained from matured adult WT and *Rnf8*-KO mice were homogenized in RIPA buffer (50 mM Tris-HCl pH 7.5, 150 mM NaCl, 0.1% SDS, 1% Triton X-100, 1% sodium deoxycholate) containing cOmplete Protease Inhibitor Cocktail (Roche, 11697498001); then, the homogenate was incubated on ice for 30 min. After DNA fragmentation by sonication and subsequent centrifugation at 10,000×*g* at 4°C, the supernatant was transferred to a new tube before total protein concentration was quantified via Bradford assays. Volumes of lysates containing 40 μg proteins were separated by electrophoresis on 10% SDS-PAGE gels. Then, the proteins were transferred onto a PVDF membrane (EMD Millipore, IPVH00010). The membranes were blocked with StartingBlockTM T20 (TBS) Blocking Buffer (ThermoFisher Scientific, 37543) at RT for 30 min before incubation with anti-RNF8 antibody (Proteintech, 14112-1-AP), diluted 1/2000 in Tris-buffered saline containing 0.1% Tween 20 detergent (TBST), at 4°C overnight. On the next day, after washing three times in TBST, 5 min per wash, the blot was incubated with VeriBlot for IP Detection Reagent (HRP) (Abcam, ab131366), diluted 1/5000 in TBST at RT, for 1 h. The blot was washed three times in TBST, 5 min per wash, before incubation in Immobilon Western Chemiluminescent HRP Substrate (EMD Millipore, WBKLS0500) at RT for 1 min; then, the blot was imaged using Super RX-N x-ray film (Fujifilm) and a FluorChemQ MultiImage III instrument (Alpha Innotech). To blot loading controls, the initial blot was stripped with Restore Western Blot Stripping Buffer (ThermoFisher Scientific, 21059) at RT for 20 min; then, the stripped blot was washed two times in TBST, 5 min per wash followed by blocking with StartingBlockTM T20 (TBS) Blocking Buffer at RT for 30 min, prior to incubation with anti-Lamin B1 antibody (Abcam, ab16048), diluted 1/2000 in TBST, at RT for 1 h. After washing the blot three times in TBST, 5 min per wash, the blot was incubated with VeriBlot for IP Detection Reagent (HRP), diluted 1/5000 in TBST, at RT for 1 h, and bands were visualized through the procedures described above.

## Acknowledgements

We thank R. Balhorn for information on Protamine antibodies and usage, and all members of the Namekawa and Li laboratories for discussion and helpful comments.

## Author Contributions

H.A. X.Z.L. and S.H.N. designed the research. H.A. and R.M. and Z.L. performed the experiments. H.A., R.P.A., X.Z.L. and S.H.N. wrote the manuscript. X.Z.L. and S.H.N. supervised the investigation. All authors reviewed the manuscript.

